# Evidence of a distinct *Blumeria graminis* f. sp. *tritici* pathotype structure in Australian wheat powdery mildew: implications for resistance breeding

**DOI:** 10.64898/2025.12.11.693847

**Authors:** Ancy Tony, Daniel Mullan, Huyen T.T. Phan, Francisco J. Lopez-Ruiz, Kejal N. Dodhia, Benjamin J. Saunders, Cao (Grace) Fang, Simon Ellwood, Kar-Chun Tan

**Affiliations:** The Centre for Crop and Disease Management, School of Molecular and Life Sciences, Curtin University, Bentley 6102, Australia; Intergrain Pty Ltd, Bibra Lake 6163, Australia; School of Molecular and Life Sciences, Curtin University, Bentley 6102, Australia

**Keywords:** *Blumeria graminis* f. sp. *tritici*, powdery mildew resistance, pathotype structure, virulence diversity, resistance breeding

## Abstract

The obligate biotrophic fungus *Blumeria graminis* f. sp. *tritici* (*Bgt*) is the causal agent of wheat powdery mildew. To investigate virulence diversity, 30 Australian *Bgt* isolates collected between 2020 and 2024 were screened against a panel of 24 wheat lines carrying defined resistance (*R*) genes. Pathotyping analysis indicated that the Australian *Bgt* population is diverse in its virulence profile. Non-metric multidimensional scaling analysis of disease phenotyping revealed evidence of six distinct *Bgt* pathotype groups. In addition, we identified wheat varieties carrying defined *R* genes and their combinations that remain effective against the contemporary Australian *Bgt*. Notably, wheat cvs Khapli and Tabasco, possessing *Pm4a* and *Pm2*, respectively, in addition to unknown *R* gene(s), were highly resistant to all the isolates tested. Furthermore, elite European varieties that possess durable resistance, such as Alchemy and Cordiale, display complete to moderate resistance against all selected isolates representing the Australian pathotype groups.

---

The obligate fungal pathogen *Blumeria graminis* f. sp. *tritici* (*Bgt*) causes wheat powdery mildew (WPM) that results in typical yield losses of 10–15% and up to 50% under favourable conditions (Rana et al. 2022). In Australia, around 12.6 million hectares are sown with wheat annually, providing a large breeding ground for *Bgt* (Department of Agriculture 2025). The fungus develops haustoria in epidermal cells to extract nutrients and reproduces asexually through airborne conidia, while chasmothecia are formed sexually under stress or at the end of the season (Spanu 2014; Wicker et al. 2013). The deployment of genetic resistance in wheat is an environmentally sustainable strategy to minimize fungicide dependence and mitigate associated ecological impacts. Resistance to WPM comprises race-specific resistance, mediated by major WPM resistance (*R/Pm*) genes that induce hypersensitive responses, and adult-plant or partial resistance, mostly a slow-mildewing form associated with broad-spectrum and durable protection (Alam et al. 2011; Ge et al. 2021). To date, over 100 *Pm* genes and allelic variants (*Pm1*-*Pm71*) conferring major and minor resistance to *Bgt* have been characterized in wheat, but only a small subset of these have been cloned (Kunz et al. 2025; Li et al. 2025). To achieve durable resistance, resistance-associated quantitative trait loci must exhibit consistent effects across multiple years and diverse environmental conditions, while maintaining stable expression throughout plant development (Kang et al. 2020).

Most wheat cultivars currently grown in Australia exhibit partial to complete susceptibility to *Bgt* (Shackley et al. 2025). The limited presence of widespread resistance to WPM in current germplasm highlights the urgent need for improvement, especially in light of proliferating fungicide resistance (Ireland et al. 2022). Demethylation inhibitor (DMI) and quinone-outside inhibitor resistance have recently become widespread in Eastern Australia, and DMI-resistance mutations have also been detected in Western Australia, as reported by the Australian Fungicide Resistance Extension Network (2025). This underscores declining fungicide efficacy against WPM (GRDC 2025; Lopez-Ruiz et al. 2023).

Race-specific resistance underlies the existence of pathotypes, or variants with a distinct virulence pattern on host genotypes (Thrall et al. 2001). Monitoring WPM pathotype structure across Australian agro-ecological zones is crucial for tracking virulence variation and guiding effective, region-specific resistance deployment in geographically isolated growing regions, as different commercial wheat varieties adopted by Australian regions affect *Bgt* adaption to varietal differences (Brown and Harris 2020; Shackley et al. 2023). The aims of this study were to determine the pathotypes present in Australia, analyse virulence differences among geographic locations, and determine the resistance status of selected sources of *Pm* genes against the current Australian *Bgt* population.

Thirty *Bgt* isolates were collected from major wheat-growing regions across Australia between 2020 and 2024 (Supplementary Table 1 and Supplementary Figure 1). Monoconidial isolates were established and maintained on the susceptible wheat cv.

Trojan using a high-throughput detached leaf assay (DLA) method for infection studies as previously described (Lopez-Ruiz et al. 2023; Tucker et al. 2013). To assess pathogenicity, a panel of 24 wheat varieties carrying different combinations of defined *Pm* genes sourced from the Australian Grains Genebank (AGG) was evaluated to identify effective *Pm* genes and determine any evidence of local *Bgt* pathotypes (Supplementary Table 2). Selected Australian commercial wheat lines were also evaluated against WPM. These varieties are Brumby (rated as resistant except to a pathotype), DS Pascal (resistant to moderately resistant), Rockstar (Moderately susceptible to susceptible) and Trojan (susceptible) (Shackley 2022). All wheat varieties were sown in individual pots with three replicates and grown in an environment-controlled growth chamber. Seven-day-old seedling leaf sections were then excised and placed onto benzimidazole agar plates. Isolate inoculation was performed using an inoculation tower and plates were incubated at 20°C under a 12:12 h light:dark photoperiod as previously described (Tucker et al. 2013). Disease severity was scored at seven days past infection based on a five-point (0-4) infection type (IT) scale (Supplementary Figure 2). 0 and 1 were considered highly resistant (R), 2 moderately resistant (MR), 3 moderately susceptible (MS), and 4 highly susceptible(S) (Li et al. 2019).

We used a heatmap approach to visualise the virulence profile of all 30 *Bgt* isolates and observed significant variations in the virulence profiles (Figure 1). Isolates 2023-060 (SA), 22PMWR15 (WA) and 20PMFRG01 (WA) were highly virulent on the wheat panel, whereas 2024PM121 (WA), 2023-046 (SA), 2024PM111 FT168 (QLD) and 2024PM108 (WA) were the least. Wheat cvs Khapli and Tabasco, both carrying an unknown *Pm* gene as well as *Pm4a* and *Pm2*, respectively, exhibited complete resistance to all the isolates tested. The Australian wheat cv Brumby possesses the highest disease resistance rating against WPM. However, resistance was overcome by the highly virulent isolate 2023-060 from a field in Malinong (SA), as was previously observed in another field location elsewhere in SA (Shackley 2022). Despite this, our study indicates that wheat cvs Khapli and Tabasco possess highly effective resistance genes against isolate 2023-060 (Supplementary Table 3). Wheat cvs Mich.Amber/8Cc, Seri 82, and Sonora/8Cc possess *Pm3f, Pm8* and *Pm3c*, respectively, and displayed the lowest levels of resistance. Thus, these *Pm* genes are ineffective against the current Australian *Bgt* population. It was observed that cv Ulka/8Cc, possessing *Pm2* (Briggle 1969), shows a mixed resistance response to the isolate panel, whereas Tabasco, also possessing *Pm2* (Wu et al. 2023), is resistant to all isolates. This strongly suggests that cv Tabasco carries an unknown *Pm* gene in addition to *Pm2*. Likewise, although cvs Kavkaz and Seri 82 both carry *Pm8* (Hurni et al. 2013; Villareal et al. 1998), their contrasting responses suggest the presence of additional resistance in Kavkaz.

**Figure 1.**
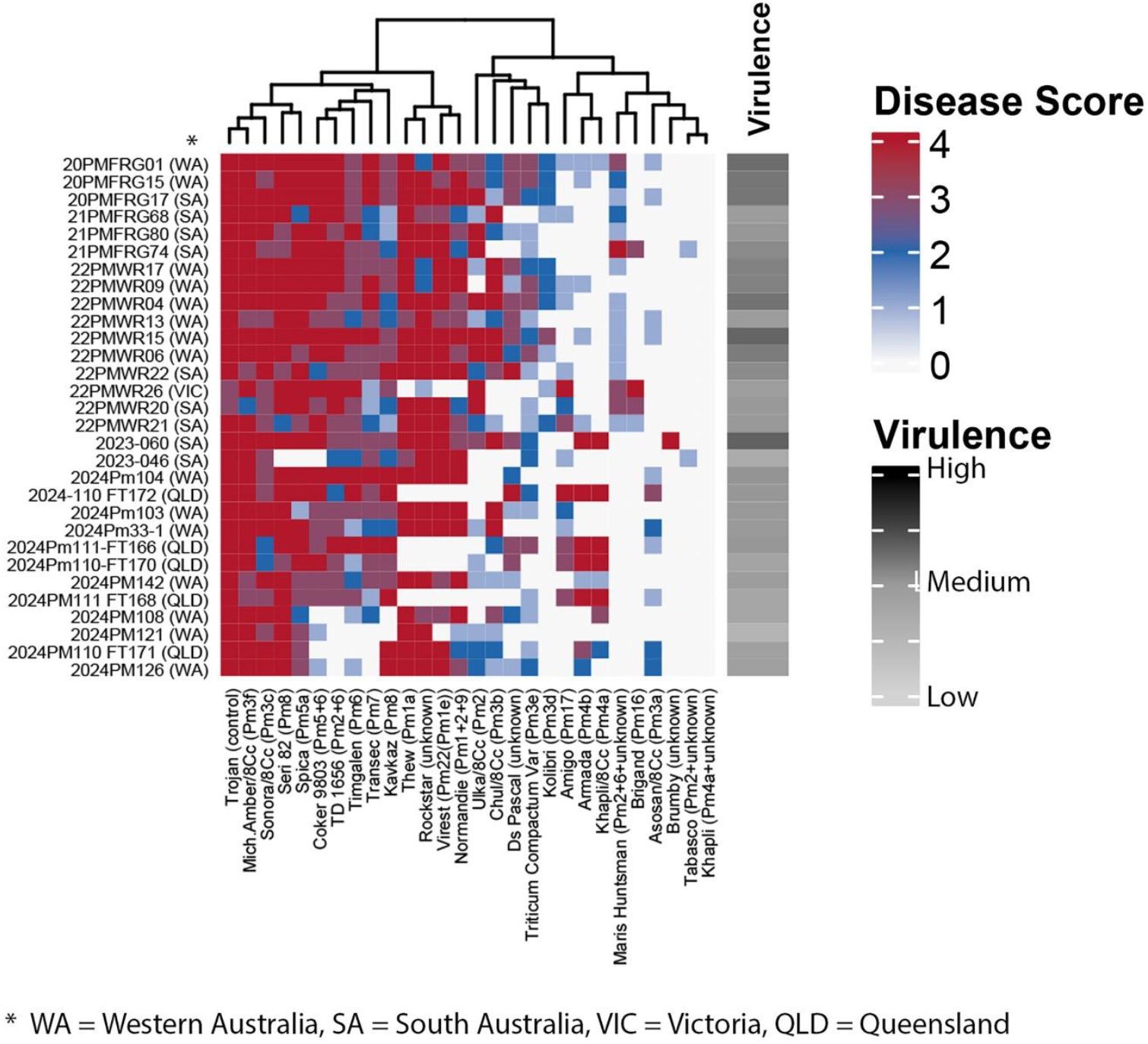
Virulence profiling of Australian *Bgt* isolates on differential and commercial wheat lines. The heatmap represents the virulence spectrum of *Bgt* isolates (*x*-axis) tested on a set of differential wheat lines (*y*-axis). The heatmap was generated using *ComplexHeatmap* package (ver.2.22.0) in R (ver.4.4.1) (Gu et al. 2016). Compiled using Adobe Photoshop (ver. 25.12.3). Detailed disease scores are provided in Supplementary Table 3.

A two-dimensional non-metric multidimensional scaling (2D nMDS) analysis was performed to assess whether virulence variability across the panel reflected any underlying population structure. The disease severity score data were exported and analysed in PRIMER 7 (ver. 7.0.24), and a 2D nMDS ordination plot was constructed based on the Euclidean distance. A similarity profile then used to visualise the *Bgt* clustering pattern that is indicative of pathotypes (M. J. Anderson 2008) (PRIMER 7 PERMANOVA+ add-on, PRIMER-E Ltd). 2D nMDS ordination identified six distinct pathotype groups (Figure 2). Correlation vectors demonstrate the contribution of specific host *R* genes to the observed variation (Figure 2A; Supplementary Figure 3). Thus, the longer the vector, the greater its influence in differentiating isolates and their orientation reflects the direction of increasing susceptibility to those isolates. cv Khapli, which exhibited an IT of 0 against all isolates, is positioned near the centre of the correlation vectors due to the lack of variation in its effectiveness. Wheat cvs Virest, Thew, Normandie and Rockstar exhibited similar susceptibility patterns and were the major contributors to pathotype IV formation. Brumby was susceptible only to isolate 2023-060, and it aligns with pathotype V, which carries the corresponding virulence. Wheat cvs Khapli/8Cc and Armada displayed related *Bgt* susceptibility profiles, with both showing pronounced susceptibility to QLD-derived isolates that predominantly define pathotype I. Wheat cvs TD 1656, Timgalen and Coker 9803 exhibited similar *Bgt* susceptibility profiles and contributed to the clustering of isolate 22PMWR26 (VIC), together with the variety Brigand, which showed complete susceptibility exclusively to this isolate. Out of these pathotypes, III and V, each carrying a single isolate, were collected in 2023 from South Australia. However, III contains the second least virulent isolate (2023-046), and the most virulent isolate (2023-060) is assigned to pathotype V. To assess the geographical variation of these pathotypes, a 2D nMDS plot was generated to determine if there is any correlation between state collection location and pathotype. Clustering of isolates by location is difficult to conclude due to insufficient isolates collected from VIC and among pathotype groups (Figure 2B). We then examined for temporal variation (Figure 2C). This indicated pathotypes I and II were comprised solely of isolates from 2024. Some isolates collected from 2024 are also found in pathotype IV, which consists of isolates collected in 2020, 2021, 2022 and 2024 (Figure 2C). However, more isolates are needed to conclusively determine whether there is a spatial-temporal effect on virulence.

**Figure 2.**
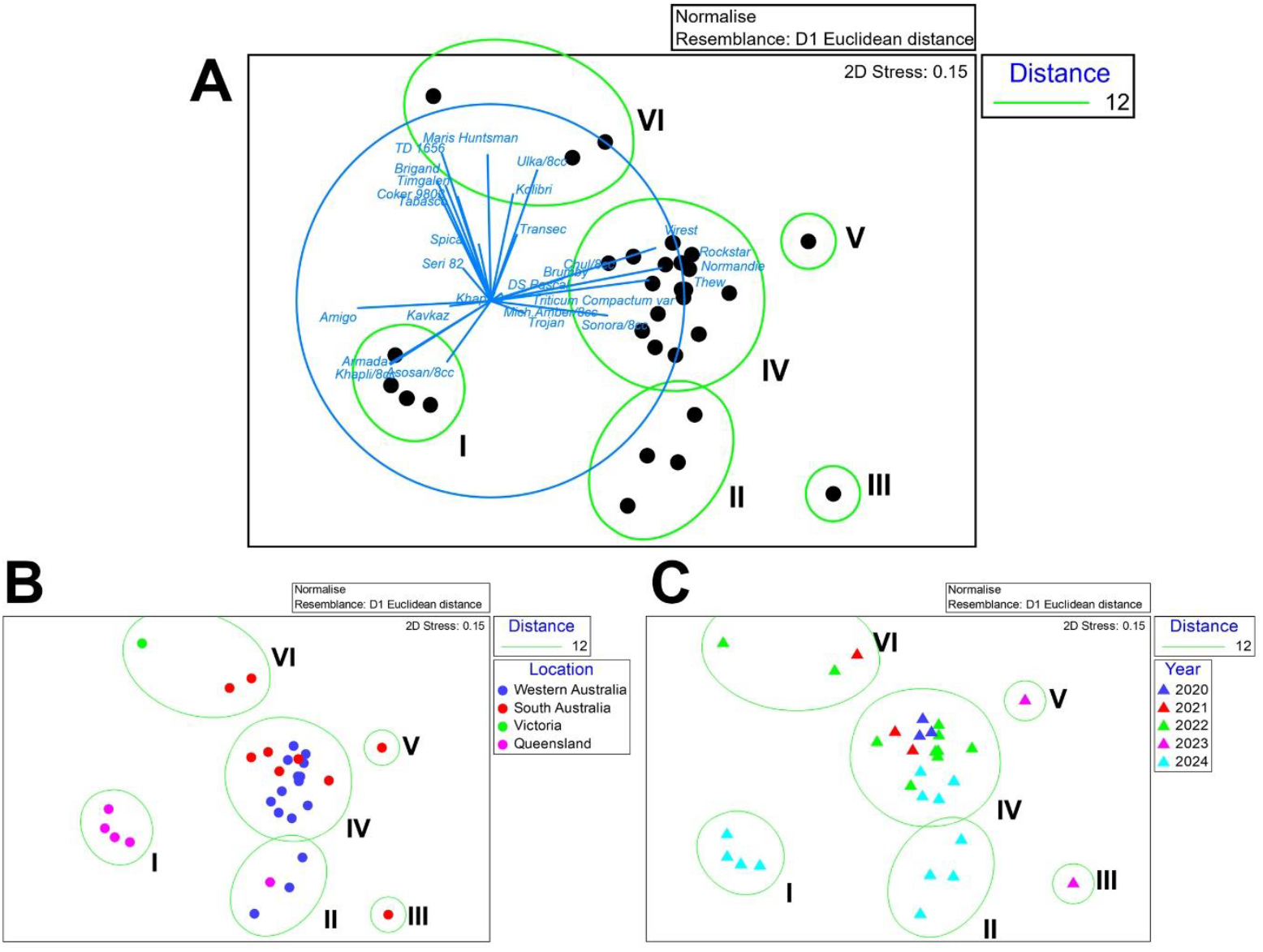
2D nMDS scaling ordination plot reveals distinct Australian *Bgt* pathotypes. A Euclidean distance similarity matrix was calculated based on the pairwise virulence profile of 30 *Bgt* isolates and used to generate nMDS coordinates of each sample and was overlaid with the hierarchical clustering analysis of these isolates, denoted as green ellipses or as groups in roman numerals. **A**, Each isolate, denoted by black dots and is overlaid with the correlation vectors calculated based on the Pearson correlation, denoted as blue lines with wheat varieties. **B**, Isolate plotted based on collection location. **C**, Isolate plotted based on the year of collection.

Single major *R* genes are often rapidly overcome by new virulent pathogen races due to strong selection pressure, whereas durable resistance remains effective over time and across environments. Durable polygenic resistance provides complete to partial protection against a wider range of pathotypes by lowering the selection pressure (Burdon et al. 2014). In the United Kingdom, Alchemy, Cordiale, and Einstein demonstrated long-lasting resistance against the local *Bgt* population over a 10-year disease trial survey (Brown 2015). These varieties were screened to assess their effectiveness against the Australian *Bgt* population by selecting representative isolates from each pathotyping group and screened using seedling DLAs. Pathotype VI was excluded due to loss of isolates. Wheat cvs Alchemy and Cordiale were resistant to all pathotypes, whereas Einstein displayed moderate resistance across all pathotypes (Table 1). Even though pathotype V contains the most virulent isolate, all three elite European cultivars showed moderate to strong resistance. This suggests the potential effectiveness of these varieties in providing long-term broad-spectrum WPM resistance in Australia. However, further testing over several years across diverse geographical and environmental conditions is required.

**Table 1.** Disease phenotypes of European durable WPM resistant wheat lines to Australian *Bgt*. Each column contains representative isolates of different pathotypes, and each row represents wheat lines. Wheat cv. Khapli (resistant) and Trojan (susceptible) were used as controls and scored/rated based on 0-4 numerical scale as described above (Li et al. 2019).

|  | Pathotype I | Pathotype II | Pathotype III | Pathotype IV | Pathotype V |
| --- | --- | --- | --- | --- | --- |
|  | 2024PM110 | 2024PM126 | 2023-046 | 20PMFRG01 | 2023-060 |
|  | 172 |  |  |  |  |
| **Alchemy** | R | R | R | R | R |
| **Cordiale** | R | R | R | R | R |
| **Einstein** | MR | MR | R | MR | MR |
| **Khapli** | R | R | R | R | R |
| **Trojan** | S | S | S | S | S |

Knowledge on Australian *Bgt* aggressiveness is limited, with only two existing published data focused exclusively on WA isolates. For instance, Golzar et al.(2016) pathotyped cumulative pools of WA *Bgt* isolates collected between 2011 and 2014, whereas Kloppe et al. (2022) characterised 15 WA isolates collected in 2016. We then compared these earlier findings with those from the contemporary Australian isolates analysed in this study (Table 2). For statistical analysis, the disease scores from both studies were converted to a 0–4 numerical scale and assigned infection types following the method described by Li et al. (2019). Comparisons of WA isolates revealed that *Pm3b, Pm3e, Pm1+2+9*, and *Pm2+6* exhibited a loss of effective resistance over time. In contrast, *Pm4b* and *Pm17* demonstrated renewed effectiveness against the WA *Bgt* isolates examined in this study. *Pm3a, Pm4a* and the *Pm4a+unknown R* gene (from cvs. Asosan/8Cc, Khapli/8Cc and Khapli, respectively) maintained their effectiveness over time in WA, with *Pm3a* and *Pm4a+unknown* also remaining effective against *Bgt* isolates from other Australian states. The effectiveness of these identified *Pm* genes require further experimental validation to ensure that WPM resistance in varieties that carry them is not due to unknown background resistance. However, Golzar et al.(2016) observed that Khapli demonstrated adult plant WPM resistance.

**Table 2.** Comparative analysis of previous pathotyping studies on Australian *Bgt* with this study. ‘NA’ indicates *Pm* genes not examined. Seedling infection scores from these studies were converted to a 0 to 4 numeric scale for statistical comparisons (Li et al. 2019). ^*^ Indicates that the wheat varieties used for the particular *Pm* genes differ among studies.

|  | (Golzar<br>et al.<br>2016) | (Kloppe<br>et al.<br>2022) | This study |  |  |  |
| --- | --- | --- | --- | --- | --- | --- |
| Region | WA | WA | WA | SA | QLD | VIC |
| R gene/sampling<br>year | 2011-2014 | 2016 | 2020-2024 |  |  |  |
| *Pm1a* * | S | S | S | S | R | R |
| *Pm2* | NA | R | MR | MR | R | S |
| *Pm3a* | R | R | R | R | R | R |
| *Pm3b* | NA | MR | MS | MR | R | R |
| *Pm3c* | S | NA | S | MS | MS | MS |
| *Pm3e* * | R | NA | MR | R | R | R |
| *Pm4a* | R | R | R | R | S | R |
| *Pm4a+unknown* | R | NA | R | R | R | R |
| *Pm4b* * | S | R | R | R | S | R |
| *Pm5a* | S | NA | MS | MS | S | S |
| *Pm6* * | S | S | MR | MS | MR | S |
| *Pm7* | MS | NA | MS | MR | MR | R |
| *Pm8* * | S | MR | MR | MR | S | MS |
| *Pm17* | S | R | R | R | MS | S |
| *Pm1+2+9* | R | NA | S | MS | R | R |
| *Pm2+6* | R | NA | MS | MS | MR | S |

Since 2020, severe WPM outbreaks have occurred across the eastern states of Australia, causing yield losses of up to 50% (Ellwood et al. 2024). Strategic use of resistance genes is only effective when supported by detailed characterisation of *Bgt* virulence and pathotype diversity (Czembor et al. 2014; El-Shamy and Mohamed 2022; Lalošević et al. 2021; Parks et al. 2008). Since the deployment of a major *R* gene often leads to boom-and-bust disease dynamics driven by the emergence of new pathogen races (Pink and Puddephat 1999). This underscores the need for continued monitoring of pathogen diversity and resistance gene efficacy to guide region-specific disease management (Zou et al. 2023).

This study represents the first reported simultaneous pathotyping of *Bgt* across a broad coverage of major wheat-growing regions in Australia, resulting in the identification of multiple virulences and six distinct pathotype groups. As many of the most popular wheat varieties in Australia are susceptible to WPM, this suggests a history of exposing *R*-genes in the field, many of which have lost their effectiveness. Despite this, wheat cvs Tabasco and Khapli exhibit complete resistance to the tested Australian *Bgt* population and represent valuable sources of resistance that can be deployed by the cereal industry into existing parental varieties. In addition to major *R* genes, the use of durable broad-spectrum resistance from Alchemy and Cordiale to minimise the impact of WPM in Australia highlights the potential of tapping into international germplasm as a resource for WPM resistance breeding in Australia. Findings from this study support targeted deployment and stacking of effective *R* genes and, ideally, implementing broad-spectrum WPM resistance. Materials collected for the current research can be used to study WPM evolution, reveal their origins, and identify their virulence factors. Furthermore, we hypothesise novel *Pm* genes effective against Australian *Bgt* are present and remain undiscovered in the differential wheat panel. Efforts are underway to locate these unknown genes and to assess their efficacy in disease control.

## Acknowledgements

We thank the AGG (Horsham, Australia) for the provision of wheat germplasm used in this study. Sam Trengove (Trengove Consulting), Hossein Golzar (DPIRD), Andrea Hills (DPIRD) and Ciara Beard (DPIRD) for the provision of *Bgt* samples. Julie Lawrence (Curtin University) for valuable technical assistance. Kul Adhikari (Curtin University), Steven Chang (Curtin University), Marta Lopez-Ruiz (Curtin University), and Nirmala Sharma (Curtin University) for isolate maintenance.

## Funding

Support was provided by Intergrain Pty Ltd (Grant CTR18532) and the Centre for Crop and Disease Management, a co-investment between the Grains

Research and Development Corporation and Curtin University (Grant CUR1403-002BLX).

## Notes

### Competing Interest Statement

The authors have declared no competing interest.

### Summary of Updates

The supplemental files have been updated. Comments present in the initial submission have been removed in the revised version. The positions of the tables have been rearranged in the main manuscript.

