## Supplementary materials for "Evidence of a distinct *Blumeria graminis* f. sp. *tritici* pathotype structure in Australian wheat powdery mildew: implications for resistance breeding"

**Supplementary Table 1:** A list of *Bgt* isolates used in this study. State locations are provided in Supplementary Figure 1.

| ***Bgt* isolate** | **Location** | **State** | **Year** | **Wheat variety** |
|---|---|---|---|---|
| 20PMFRG01 | Cranbrook | Western Australia | 2020 | Unknown |
| 20PMFRG15 | Esperance | Western Australia | 2020 | Unknown |
| 20PMFRG17 | Bute | South Australia | 2020 | Unknown |
| 21PMFRG68 | Unknown | South Australia | 2021 | Unknown |
| 21PMFRG80 | Unknown | South Australia | 2021 | Unknown |
| 21PMFRG74 | Unknown | South Australia | 2021 | Unknown |
| 22PMWR17 | Manjimup | Western Australia | 2022 | Unknown |
| 22PMWR09 | Esperance | Western Australia | 2022 | Devil |
| 22PMWR04 | Geraldton | Western Australia | 2022 | Scepter |
| 22PMWR13 | South Perth | Western Australia | 2022 | Wyalkatchem |
| 22PMWR15 | Marchagee | Western Australia | 2022 | Unknown |
| 22PMWR06 | Wongan Hill | Western Australia | 2022 | Catapult |
| 22PMWR22 | Unknown | South Australia | 2022 | Unknown |
| 22PMWR26 | Unknown | Victoria | 2022 | Unknown |
| 22PMWR20 | Unknown | South Australia | 2022 | Unknown |
| 22PMWR21 | Unknown | South Australia | 2022 | Unknown |
| 2023-060 | Malinong | South Australia | 2023 | Brumby |
| 2023-046 | Port Broughton | South Australia | 2023 | Scepter |
| 2024Pm104 | Mount Barker | Western Australia | 2024 | Scepter |
| 2024-110 FT172 | Pampas | Queensland | 2024 | Leverage |
| 2024Pm103 | Scotts Brook | Western Australia | 2024 | Rockstar |
| 2024Pm33-1 | Lake Grace | Western Australia | 2024 | Unknown |
| 2024Pm111FT166 | Pampas | Queensland | 2024 | Sunflex |
| 2024Pm110FT170 | Pampas | Queensland | 2024 | Sunflex |
| 2024PM142 | Kendenup | Western Australia | 2024 | Kinsei |
| 2024PM111 FT168 | Pampas | Queensland | 2024 | Leverage |
| 2024PM108 | Cascade | Western Australia | 2024 | Scepter |
| 2024PM121 | Kendenup | Western Australia | 2024 | Kinsei |
| 2024PM110 FT171 | Pampas | Queensland | 2024 | Sunflex |
| 2024PM126 | Gibson | Western Australia | 2024 | Unknown |

**Supplementary Figure 1:** Map depicting the originating states of *Bgt* isolates. The size of the circle represents the total number of isolates collected from each state (denoted as n). The segment size and colour coding of the pie chart represent the number of isolates collected each year. Map generated using R Statistical Software (R Core Team, 2024) and the ggplot2 package v3.5.0 (Wickham 2016). Geographic data obtained from rnaturalearth (South 2025).

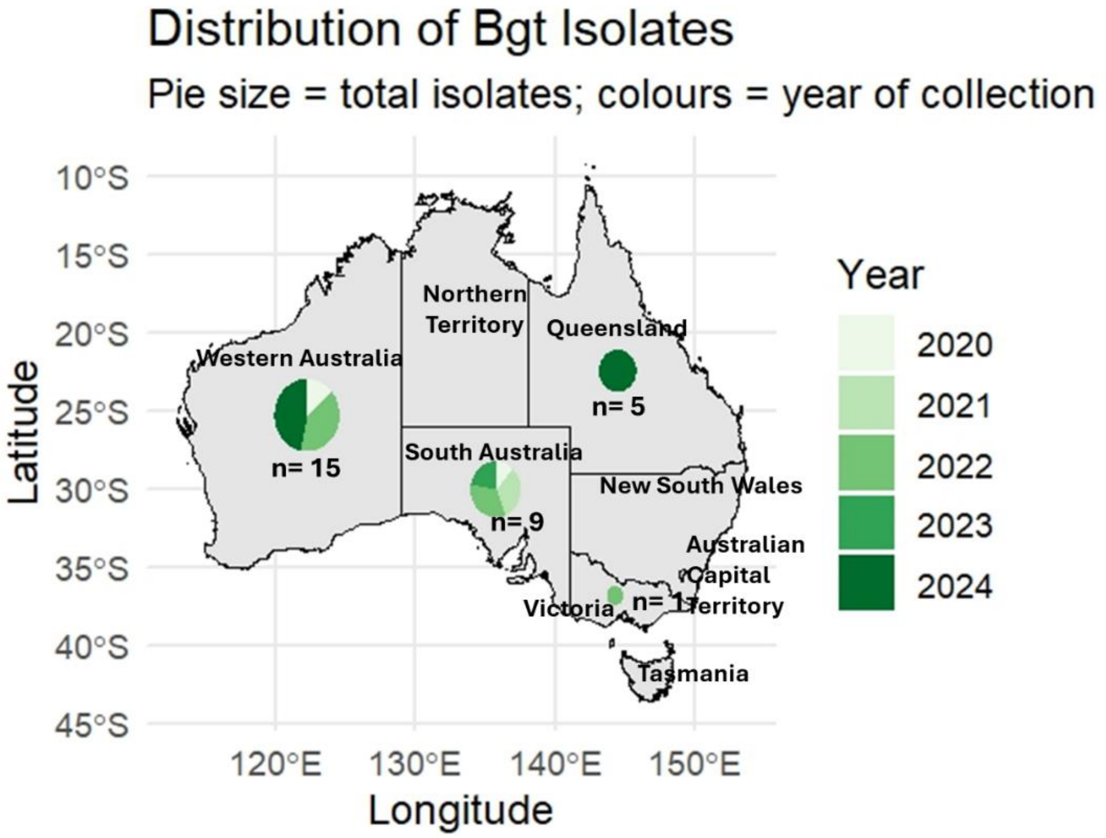

**Supplementary Table 2:** Differential wheat lines carrying *Pm* genes used in this study. *Pm?* = undiscovered *R* genes, 8Cc = near-isogenic lines derived from the recurrent parent Chancellor, NBS-LRR = Nucleotide-Binding Site (NBS) and Leucine-Rich Repeat (LRR) domains, STKs = serine/threonine protein kinase, NA = no information available

| **Wheat lines** | ***R* gene(s) ID** | **Chromosome location** | ***R* gene class** | **References** |
|---|---|---|---|---|
| Spica | *Pm5a* | 7BL | NBS-LRR | (Law and Wolfe 1966; Xie et al. 2020) |
| Thew | *Pm1a* | 7AL | NBS-LRR | (Liu et al. 2017; Sears and Briggle 1969) |
| Transec | *Pm7* | T4BS.4BL-5RL | NA | (Heun and Friebe 1990) |
| Kolibri | *Pm3d* | 1AS | NBS-LRR | (Srichumpa et al. 2005; Zeller et al. 1993b) |
| Normandie | *Pm1+2+9* | 7AL,5DS, | NBS-LRR | (Schneider et al. 1991) |
| TD 1656 | *Pm2+6* | 5DS,2BL | NBS-LRR | (Golzar et al. 2016) |
| Timgalen | *Pm6* | 2BL | NBS-LRR | (Bennett and van Kints 1983; Chen et al. 2016; Tao et al. 2000) |
| Virest | *Pm22(Pm1e)* | 7AL | NA | (Singrün et al. 2003) |
| Ulka/8Cc | *Pm2* | 5DS | NBS-LRR | (Briggle 1969; Manser et al. 2021) |
| Mich.Amber/8Cc | *Pm3f* | 1AS | NBS-LRR | (Briggle 1969; Srichumpa et al. 2005) |
| Asosan/8Cc | *Pm3a* | 1AS | NBS-LRR | (Briggle 1969; Srichumpa et al. 2005) |
| Chul/8Cc | *Pm3b* | 1AS | NBS-LRR | (Briggle 1969; Srichumpa et al. 2005) |
| Sonora/8Cc | *Pm3c* | 1AS | NBS-LRR | (Briggle 1969; Srichumpa et al. 2005) |
| Khapli/8Cc | *Pm4a* | 2AL | STKs | (Briggle 1966; Huang et al. 1997) |
| Kavkaz | *Pm8* | T1RS.1BL | NBS-LRR | (Hurni et al. 2013) |
| Maris Huntsman | *Pm2+6+Pm?* | 5DS,2BL | NBS-LRR | (Kowalczyk et al. 1998) |
| Amigo | *Pm17* | 1RS.1AL | NBS-LRR | (Hsam and Zeller 1997) |
| Brigand | *Pm16* | 4A | NA | (Wu et al. 2021) |
| Armada | *Pm4b* | 2AL | STKs | (Sánchez-Martín et al. 2021; Zeller et al. 1993a) |
| Seri 82 | *Pm8* | 1RS.1BL | NBS-LRR | (Hurni et al. 2013; Villareal et al. 1998) |
| *Triticum compactum var* | *Pm3e* | 1AS | NBS-LRR | (Srichumpa et al. 2005; Wu et al. 2021) |

| | | | | |
|---|---|---|---|---|
| Coker 9803 | *Pm5+6* | 7BL,2BL | NBS-LRR | (Wu et al. 2021) |
| Tabasco | *Pm2* | 5DS | NBS-LRR | (Manser et al. 2021; Wu et al. 2023) |
| Khapli | *Pm4a+Pm?* | 2AL | STKs | (Vida et al. 2002) |

**Supplementary Figure 2:** *Bgt* infection types of the 0-4 wheat numerical scoring scale. 0= no visible symptoms; 1= low fungal development without sporulation; 2= moderate mycelial growth with slight sporulation; 3= dense mycelial growth with high sporulation and less colonisation; 4= large colony growth with extensive sporulation. 0 and 1 are rated as highly resistant (R), 2 moderately resistant (MR), 3 moderately susceptible (MS), and 4 highly susceptible (S) (Li et al. 2019).

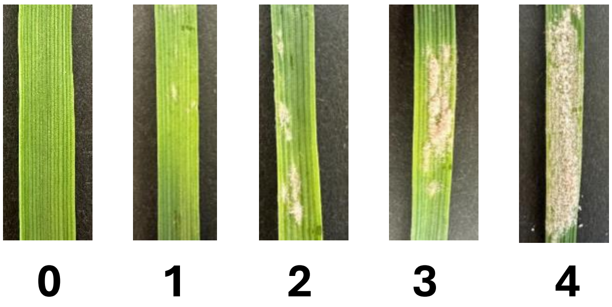

**Supplementary Table 3:** Wheat varietal resistance and virulence profiles of Australian *B. graminis* isolates collected between 2020 and 2024. Abbreviations are WA = Western Australia, SA = South Australia, VIC = Victoria, NSW/QLD = New South Wales/Queensland, - = no data available, and +*Pm?* = additional unknown *R* gene or genes. *Pm22(Pm1e)* = *Pm22* in cultivar Virest is a member of the complex *Pm1* locus in common wheat. Disease score rating for Rockstar, DS Pascal, Brumby and Trojan are moderately susceptible to susceptible (MSS), resistant-moderately resistant (RMR), resistant (R) and susceptible (S) (Shackley 2022). Colour scale ranges from green (resistant wheat) to red (susceptible wheat) (legend provided). Dark to light purple denotes the highest virulent to the lowest virulent isolate, respectively (legend provided).

| Differential Lines | *R* gene(s) ID | 20PMFRG01 | 20PMFRG15 | 20PMFRG17 | 21PMFRG68 | 21PMFRG80 | 21PMFRG74 | 22PMWR17 | 22PMWR09 | 22PMWR04 | 22PMWR13 | 22PMWR15 | 22PMWR06 | 22PMWR22 | 22PMWR26 | 22PMWR20 | 22PMWR21 | 2023-060 | 2023-046 | 2024Pm104 | 2024-110 FT172 | 2024Pm103 | 2024Pm33-1 | 2024Pm111-FT166 | 2024Pm110-FT170 | 2024PM142 | 2024PM111 FT168 | 2024PM108 | 2024PM121 | 2024PM110 FT171 | 2024PM126 | Total effective resistance score |
|---|---|---|---|---|---|---|---|---|---|---|---|---|---|---|---|---|---|---|---|---|---|---|---|---|---|---|---|---|---|---|---|---|
| | | WA | WA | SA | SA | SA | SA | WA | WA | WA | WA | WA | WA | SA | VIC | SA | SA | SA | SA | WA | QLD | WA | WA | QLD | QLD | WA | QLD | WA | WA | QLD | WA | |
| Spica | *Pm5a* | 4 | 4 | 4 | 2 | 4 | 4 | 4 | 4 | 4 | 4 | 4 | 3 | 4 | 4 | 4 | 3 | 4 | 0 | 4 | 4 | 1 | 4 | 4 | 4 | 3 | 3 | 2 | 3 | 3 | 3 | 80 |
| Thew | *Pm1a* | 4 | 4 | 4 | 4 | 4 | - | 4 | 4 | 4 | 3 | 4 | 4 | 4 | 0 | 4 | 4 | 4 | 3 | 4 | 0 | 4 | 4 | 0 | 0 | 4 | 0 | 4 | 4 | 4 | 4 | 74 |
| Transec | *Pm7* | 4 | 4 | 4 | 2 | 2 | 3 | 3 | 4 | 4 | 3 | 4 | 3 | 3 | 1 | 1 | 2 | 3 | 3 | 4 | 3 | 3 | 2 | 4 | 3 | 3 | 1 | 2 | 0 | 0 | 0 | 59 |
| Kolibri | *Pm3d* | 2 | 2 | 2 | 1 | 0 | 0 | 2 | 2 | 2 | 0 | 3 | 1 | 0 | 1 | 0 | 2 | 0 | 0 | 0 | 0 | 0 | 0 | 0 | 0 | 0 | 0 | 0 | 0 | 0 | 0 | 13 |
| Normandie | *Pm1+2+9* | 3 | 4 | 4 | 2 | 3 | 2 | 4 | 4 | 3 | 3 | 4 | 3 | 4 | 0 | 2 | 3 | 3 | 4 | 4 | 0 | 4 | 4 | 0 | 0 | 4 | 0 | 4 | 1 | 2 | 3 | 63 |
| Td 1656 | *Pm2+6* | 4 | 4 | 4 | 4 | 3 | 4 | 4 | 4 | 3 | 3 | 4 | 4 | 3 | 4 | 4 | 3 | 3 | 2 | 4 | 2 | 3 | 3 | 4 | 3 | 3 | 3 | 0 | 0 | 0 | 0 | 66 |
| Timgalen | *Pm6* | 3 | 3 | 3 | 3 | 4 | 3 | 3 | 3 | 3 | 2 | 4 | 3 | 3 | 4 | 3 | 3 | 3 | 2 | 4 | 4 | 3 | 1 | 4 | 1 | 2 | 3 | 1 | 0 | 0 | 1 | 60 |
| Virest | *Pm22(Pm1e)* | 4 | 3 | 4 | 3 | 4 | 4 | 4 | 4 | 4 | 3 | 4 | 4 | 4 | 0 | 4 | 4 | 4 | 4 | 4 | 0 | 4 | 4 | 0 | 0 | 3 | 0 | 3 | 0 | 4 | 4 | 69 |
| Ulka/8Cc | *Pm2* | 3 | 3 | 4 | 1 | 4 | 4 | 1 | 0 | 4 | 1 | 4 | - | 2 | 4 | 4 | 2 | 3 | 0 | 0 | 0 | 0 | 1 | 0 | 1 | 1 | 0 | 0 | 1 | 2 | 2 | 33 |
| Mich.Amber/8Cc | *Pm3f* | 4 | 4 | 4 | 4 | 4 | 4 | 4 | 4 | 4 | 3 | 4 | 4 | 3 | 4 | 2 | 4 | 4 | 4 | 4 | 4 | 4 | 4 | 4 | 4 | 3 | 3 | 4 | 4 | 4 | 4 | 90 |
| Asosan/8Cc | *Pm3a* | 1 | 0 | 1 | 0 | 0 | 0 | 0 | 0 | 0 | 1 | 1 | 0 | 0 | 0 | 0 | 0 | 0 | 0 | 1 | 3 | 0 | 2 | 1 | 0 | 0 | 1 | 0 | 0 | 2 | 2 | 14 |
| Chul/8Cc | *Pm3b* | 2 | 2 | 1 | 4 | 2 | 0 | 4 | 4 | 4 | 2 | 4 | 4 | 3 | 0 | 0 | 1 | 4 | 0 | 0 | 0 | 4 | 4 | 2 | 0 | 1 | 0 | 3 | 1 | 2 | 1 | 48 |
| Sonora/8Cc | *Pm3c* | 4 | 3 | 4 | 4 | 4 | 3 | 4 | 4 | 4 | 3 | 4 | 4 | 4 | 3 | 3 | 3 | 4 | 3 | 3 | 3 | 4 | 4 | 2 | 2 | 4 | 3 | 4 | 3 | 4 | 4 | 83 |

| | | | | | | | | | | | | | | | | | | | | | | | | | | | | | | | | |
|---|---|---|---|---|---|---|---|---|---|---|---|---|---|---|---|---|---|---|---|---|---|---|---|---|---|---|---|---|---|---|---|---|
| **Khapli/8Cc** | ***Pm4a*** | 1 | 0 | 0 | 0 | 0 | 0 | 0 | 0 | 0 | 0 | 0 | 0 | 0 | 0 | 0 | 0 | 4 | 0 | 0 | 4 | 0 | 0 | 4 | 4 | 1 | 4 | 3 | 0 | 2 | 0 | 26 |
| **Kavkaz** | ***Pm8*** | 3 | 3 | 3 | 1 | 1 | 2 | 3 | 3 | 2 | 2 | 3 | 3 | 4 | 3 | 2 | 1 | 3 | 2 | 4 | 4 | 3 | 2 | 4 | 3 | 3 | 4 | 0 | 0 | 4 | 4 | 66 |
| **Maris Huntsman** | ***Pm2+6+Pm?*** | 3 | 2 | 1 | 2 | 1 | 4 | 1 | 0 | 1 | 1 | 1 | 1 | 1 | 3 | 3 | 1 | 0 | 0 | 0 | 0 | 0 | 0 | 0 | 0 | 0 | 0 | 0 | 0 | 0 | 0 | 13 |
| **Amigo** | ***Pm17*** | 1 | 0 | 0 | 1 | 0 | 0 | 0 | 1 | 1 | 0 | 0 | 0 | 1 | 4 | 2 | 3 | 0 | 1 | 0 | 4 | 2 | 1 | 3 | 3 | 0 | 3 | 0 | 0 | 0 | 0 | 29 |
| **Brigand** | ***Pm16*** | 0 | 0 | 0 | 0 | 0 | 3 | 0 | 0 | 0 | 0 | 0 | 0 | 0 | 4 | 3 | 1 | 0 | 0 | 0 | 0 | 0 | 0 | 0 | 0 | 0 | 0 | 0 | 0 | 0 | 0 | 8 |
| **Armada** | ***Pm4b*** | 1 | 1 | 0 | 0 | 0 | 0 | 0 | 1 | 0 | 0 | 1 | 0 | 0 | 0 | 0 | 1 | 4 | 0 | 0 | 4 | 0 | 0 | 4 | 4 | 1 | 4 | 0 | 0 | 3 | 2 | 29 |
| **Seri 82** | ***Pm8*** | 4 | 4 | 4 | 4 | 3 | 3 | 4 | 4 | 4 | 4 | 4 | 4 | 3 | 4 | 4 | 2 | 4 | 0 | 4 | 4 | 4 | 4 | 4 | 3 | 4 | 4 | 4 | 4 | 4 | 4 | 88 |
| ***Triticum compactum var*** | ***Pm3e*** | 3 | 3 | 2 | 0 | 1 | 1 | 2 | 3 | 3 | 1 | 2 | 3 | 1 | 0 | 1 | 1 | 2 | 2 | 0 | 2 | 1 | 0 | 3 | 0 | 0 | 1 | 1 | 0 | 1 | 2 | 32 |
| **Coker 9803** | ***Pm5+6*** | 4 | 4 | 4 | 4 | 4 | 4 | 4 | 4 | 4 | 3 | 4 | 4 | 2 | 4 | 3 | 4 | 4 | 0 | 4 | 4 | 3 | 3 | 3 | 4 | 3 | 3 | 0 | 1 | 0 | 1 | 69 |
| **Tabasco** | ***Pm2+Pm?*** | 0 | 0 | 0 | 0 | 0 | 1 | 0 | 0 | 0 | 0 | 0 | 0 | 0 | 0 | 0 | 0 | 0 | 1 | 0 | 0 | 0 | 0 | 0 | 0 | 0 | 0 | 0 | 0 | 0 | 0 | 1 |
| **Khapli** | ***Pm4a+Pm?*** | 0 | 0 | 0 | 0 | 0 | 0 | 0 | 0 | 0 | 0 | 0 | 0 | 0 | 0 | 0 | 0 | 0 | 0 | 0 | 0 | 0 | 0 | 0 | 0 | 0 | 0 | 0 | 0 | 0 | 0 | 0 |
| **Rockstar** | ***MSS*** | 2 | 4 | 4 | 3 | 4 | 4 | 2 | 2 | 4 | 4 | 4 | 4 | 4 | 1 | 4 | 4 | 4 | 4 | 4 | 0 | 4 | 4 | 0 | 0 | 4 | 0 | 3 | 4 | 4 | 4 | 72 |
| **DS Pascal** | ***RMR*** | 3 | 3 | 3 | 0 | 1 | 0 | 3 | 1 | 3 | 3 | 3 | 2 | 4 | 0 | 0 | 0 | 3 | 0 | 2 | 4 | 1 | 0 | 3 | 3 | 1 | 0 | 2 | 0 | 0 | 1 | 39 |
| **Brumby** | **R** | 0 | 0 | 0 | 0 | 0 | 0 | 0 | 0 | 0 | 0 | 0 | 0 | 0 | 0 | 0 | 0 | 4 | 0 | 0 | 0 | 0 | 0 | 0 | 0 | 0 | 0 | 0 | 0 | 0 | 0 | 4 |
| **Trojan** | ***S*** | 4 | 4 | 4 | 4 | 4 | 4 | 4 | 4 | 4 | 4 | 4 | 4 | 4 | 3 | 3 | 4 | 4 | 4 | 4 | 4 | 4 | 4 | 4 | 4 | 4 | 4 | 4 | 4 | 4 | 4 | 94 |
| **Total virulence score** | | 71 | 68 | 68 | 53 | 57 | 57 | 64 | 64 | 69 | 53 | 74 | 62 | 61 | 51 | 56 | 56 | 75 | 39 | 58 | 57 | 56 | 55 | 57 | 46 | 52 | 44 | 44 | 30 | 49 | 50 | |

Resistant → Susceptible

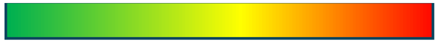

Less virulent → High virulent

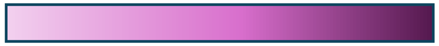

**Supplementary Figure 3:** 2D nMDS ordination of 30 Australian *Bgt* isolates based on Euclidean distances of their virulence profiles. Isolates are shown with distinct shapes and colours, with hierarchical clustering groups indicated by green ellipses (Roman numerals). Pearson correlation vectors are overlaid as blue arrows.

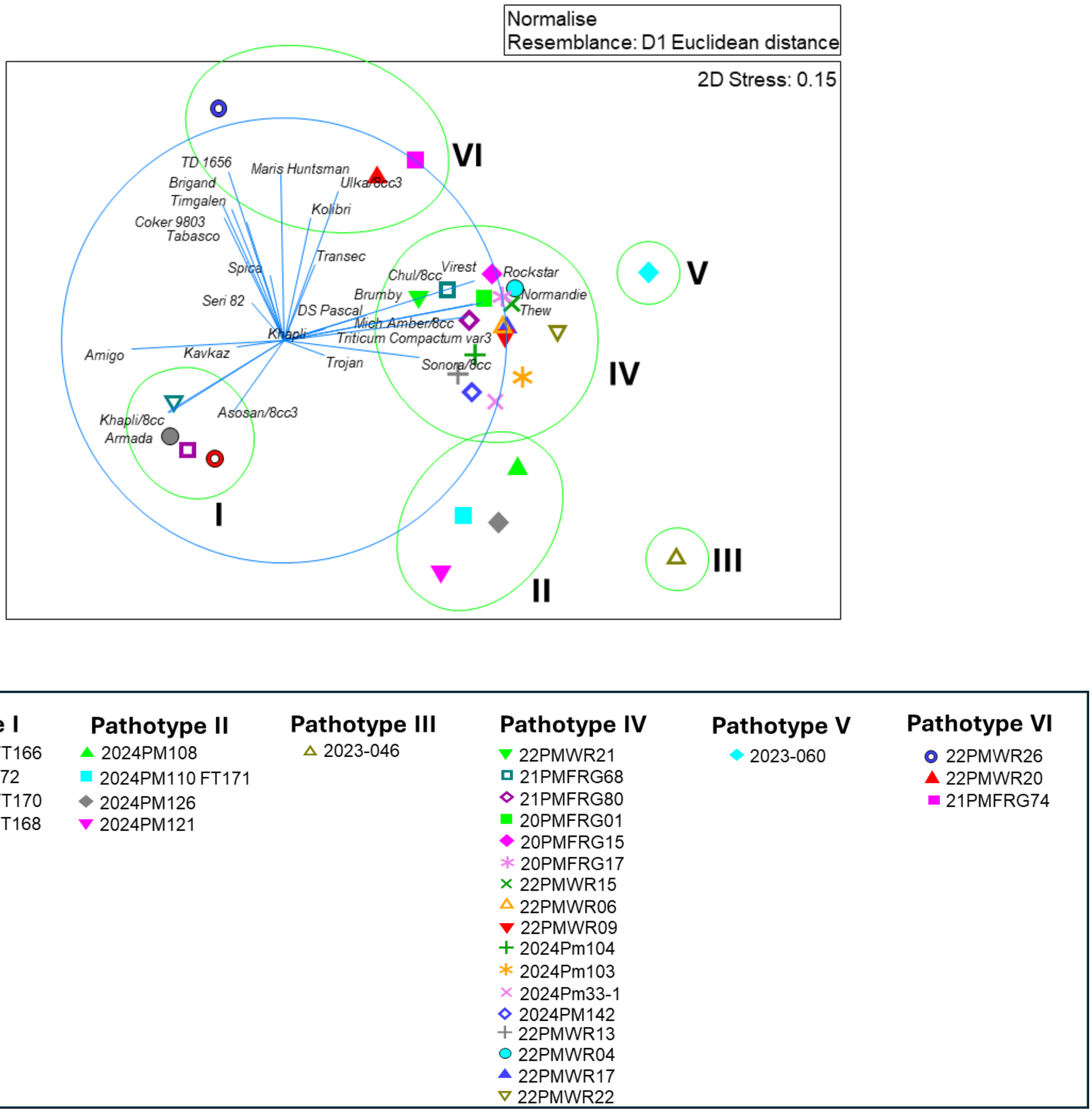
